# Beyond Life: Exploring Hemodynamic Patterns in Postmortem Mice Brains

**DOI:** 10.1101/2024.01.16.575850

**Authors:** Anton Sdobnov, Alexander Bykov, Gennadi Piavchenko, Vassiliy Tsytsarev, Igor Meglinski

## Abstract

We use Laser Speckle Contrast Imaging (LSCI) for transcranial visualization of cerebral blood flow microcirculation in mice during and after cardiac arrest. Analyzing time series of LSCI images, we observed temporal variations in blood flow distribution across the brain surface for up to several hours postmortem. Utilizing Fast Fourier Transform (FFT) analysis, we depicted the decay in blood flow oscillations and microcirculation following death. Due to the exponential drop in blood flow intensity and ensuing non-stationary conditions, Continuous Wavelet Transform (CWT) was applied to identify potential spatial or temporal synchronization patterns in cerebral hemodynamics. Additionally, we conducted Non-negative Matrix Factorization (NMF) analysis with four components to segment LSCI images, revealing temporal alterations in structural subcomponents. This integrated approach, combining LSCI, FFT, CWT and NMF, provides a comprehensive tool for understanding cerebral blood flow dynamics in mice, metaphorically capturing the ‘end of the tunnel’ experience. Results indicated a primary localization of hemodynamic activity in the olfactory bulbs postmortem, followed by minor successive relocations of blood microflows between the somatosensory and visual cortical regions via the superior sagittal sinus. The proposed approach opens avenues for further exploration into these phenomena, potentially bridging the gap between neuroscientific understanding and the longstanding mysteries surrounding consciousness and perception at the end of life.

## I. INTRODUCTION

The enigmatic experience of a ‘light at the end of the tunnel’, reported throughout history, has long intrigued both the scientific community and the public. Neurophysiological investigations following cardiac arrest have uncovered organized electrical brain signals in rats [1] and distinctive slow-wave patterns in human electroencephalograms (EEG) readings [2], phenomena often termed the ‘wave of death’. These findings only begin to unravel the complexities of brain activity during and after the death process [3], with electrical activities akin to those seen in deep sleep observed to persist even 10 minutes following the clinical declaration of [4]. Furthermore, recent revelations suggest the possibility of a post-mortem revival in brain microcirculation and cellular functions, occurring several hours after death [5]. In addition to these observations, recent years have seen increased focus on neurovascular changes during hypoxia in the terminal stage. Studies analyzing EEG data of dying patients before and after the cessation of mechanical ventilation [6, 7] have identified stimulation of gamma activity by global hypoxia in some cases. However, the underlying mechanisms and physiological significance of these observations, including those related to cerebral hemodynamics postmortem, remain largely unclear [8, 9]. This context frames our study’s aim to explore further and bridge the gap between these initial observations and a deeper understanding of postmortem brain activity.

While the causal relationship between neural activity and hemodynamics is beyond doubt, the deeper mechanisms underlying this link, especially postmortem, remain a subject of ongoing research [10–13]. Studies using Dynamic Light Scattering (DLS)-based approach and direct optical vascular imaging techniques have uncovered randomized blood microflows in various tissues of small animals for hours after death [14]. Furthermore, separate assessments of cerebral blood flow impairments caused by cardiac and respiratory arrest using multi-modal diagnostic tools have been conducted [15]. These studies reveal that cerebral hemodynamic alterations, similar to electrical brain activities, persist for several hours following respiratory arrest or cessation of blood flow. However, despite the accumulation of data, the precise mechanisms driving these cerebral hemodynamic variations and their structural organization postmortem are still not fully understood. This gap in knowledge is particularly notable when considering the substantial anatomical differences between the rodent and human brain, suggesting that the sequence of irreversible changes following oxygen deprivation may vary significantly across species.

DLS-based techniques, including LSCI, play a crucial role in brain imaging due to their unique ability to non-invasively monitor and analyze cerebral blood flow and microcirculation in real-time. These techniques are particularly significant for understanding neurovascular dynamics, aiding in the exploration of various physiological and pathological conditions within the brain [16]. LSCI, as a specific DLS-based approach, stands out for its high temporal resolution, which allows for the observation of rapid real-time hemodynamic changes, and its ease of use, making it accessible for both clinical and research settings. LSCI is a valuable tool for real-time, non-invasive monitoring of cerebral blood flow, particularly during surgeries [17], and its capability to quantitatively analyze blood flow dynamics plays an invaluable role in advancing our understanding of cerebral hemodynamics, offering a comprehensive perspective on brain function and health [18]. However, due to the complexity of the data it generates, additional statistical analysis and segmentation are often required to accurately quantify blood flow, enhance image clarity, and ensure the images are useful for medical decision-making [19].

Statistical analysis, including techniques like wavelet transform, and spectral analysis methods such as Fast Fourier Transform (FFT), enhance the interpretability of LSCI data by decomposing complex blood flow signals into simpler components [20–22]. This helps in identifying specific physiological phenomena or vascular responses, such as those occurring during reactive hyperemia. While FFT is excellent for identifying the overall frequency content of a signal, wavelet analysis [22] excels in analyzing signals where frequency components vary over time, making it a more versatile tool for detailed time-frequency analysis, especially for non-stationary signals like those often encountered in LSCI imaging.

In fact, Non-negative Matrix Factorization (NMF) [23] offers certain advantages compared to FFT and wavelet transform, particularly in the context of analyzing complex data like LSCI images. NMF tends to produce a part-based representation, where each component reflects a part or a feature of the data. This is particularly useful in image analysis, as it can lead to the identification of distinct patterns or regions within the image, which might correspond to specific physiological or anatomical structures. NMF’s strength lies in its ability to decompose data into non-negative, interpretable components, making it particularly useful for image analysis and feature extraction. This contrasts with FFT and wavelet transform, which are more focused on signal frequency and time-frequency analysis. While all FFT, wavelet analysis and NMF are valuable for the LSCI data analysis, their applications and focus in analyzing LSCI are different. FFT and wavelet analysis are particularly suited for examining temporal and spatial patterns in dynamic signals like blood flow, while NMF is more aligned with decomposing data into interpretable, static components. In the current study, we apply the Laser Speckle Contrast Imaging (LSCI) approach, specifically developed for transcranial optical vascular imaging (nTOVI) [24], to investigate cerebral hemodynamic alterations following cardiac cessation. To enhance clarity, the obtained LSCI images were processed using both FFT [25] and Continuous Wavelet Transform (CWT), specifically employing the Mexican hat wavelet [26], as well as the NMF analytical approach [27]. More specifically, we introduce the application of NMF not only for segmenting individual LSCI images into meaningful components but also for analyzing a series of LSCI images over time, thereby revealing temporal alterations in structural subcomponents.

## II. RESULTS AND DISCUSSION

The obtained results are presented in Fig 1. For the better clarity, the background area outside brain has been colored in gray.

**FIG. 1:**
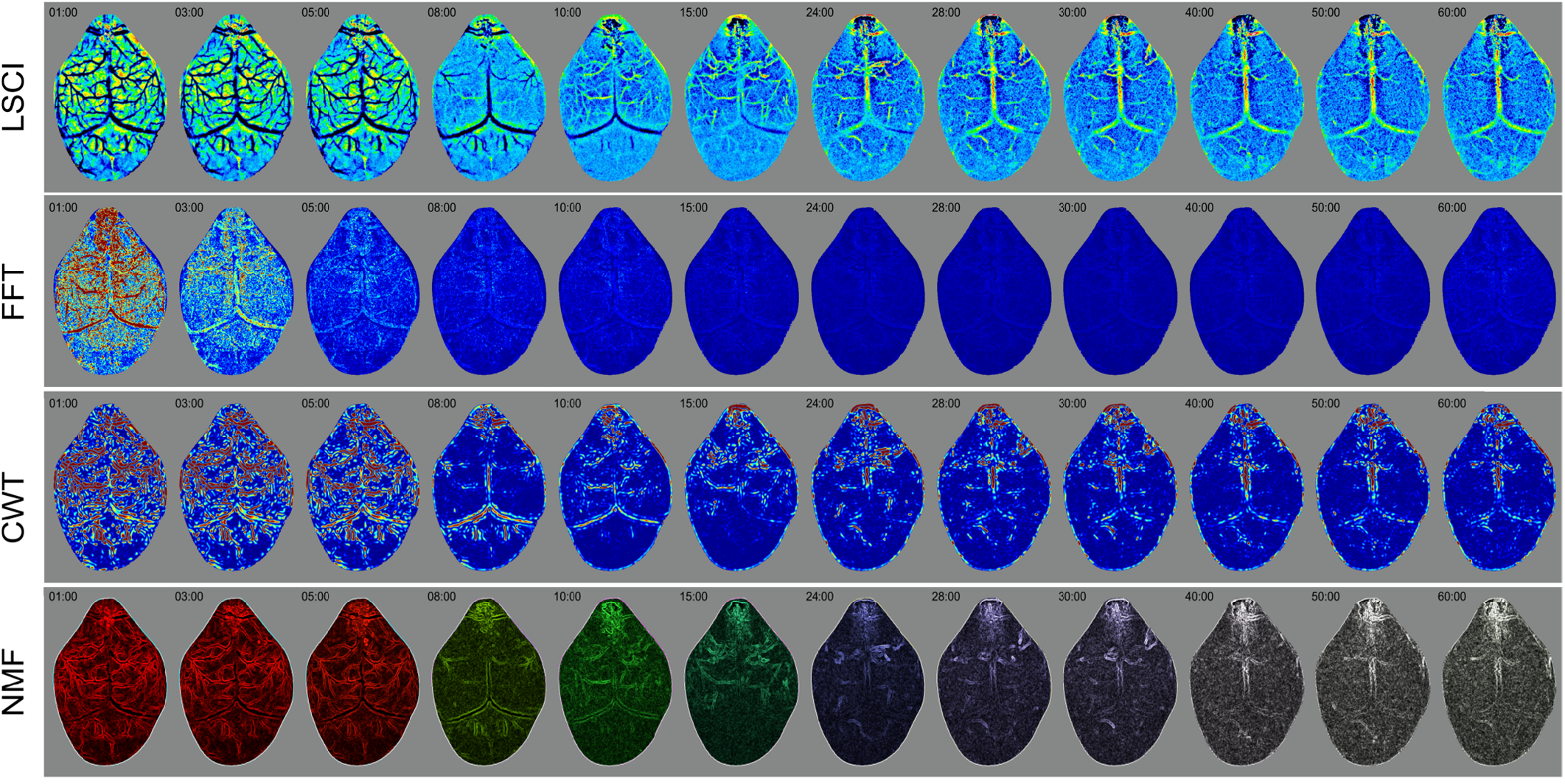
The color-coded images of mouse brain obtained, respectively, by LSCI (first line), FFT (second line), CWT (third line) and NMF (forth line) at different time intervals after the cardiac arrest. Supplementary video 1 shows full time series of LSCI, FFT, CWT and NMF. Time intervals are selected according NMF component analysis discussed below.

The time series of LSCI images reveals temporal variations in the spatial distribution of blood flow across the brain surface, as illustrated in Fig 1, line 1. Decomposition of these images using FFT captures the decay of oscillations in blood flow and microcirculation following the animal’s death, shown in Fig 1, line 2. Considering the non-stationary nature of the recorded LSCI image set, and the variability of its statistical properties over time, we applied CWT to investigate the emergence of cerebral hemodynamic patterns indicating spatial and/or temporal synchronization across the brain surface (see Fig 1, line 3). Additionally, we employed NMF for segmenting LSCI images, which aids in identifying interpretable relationships within distinct, demarcated microstructural patterns for functional evaluation, as detailed in Fig 1, line 4. To effectively characterize postmortem cerebral hemodynamics and delineate its features, we selected four NMF components, balancing stability and accuracy as referenced in previous studies [27]. These components represent distinct patterns of microstructural variance in cerebral blood microcirculation, and their temporal analysis offers profound insights into the complexity and heterogeneity of hemodynamic localization, both spatially and temporally. The NMF approach thus facilitates the identification of distinct microstructural components, enhancing our understanding of hemodynamic pattern formation in specific cortical zones and their functional implications. The color-coded representations of these four NMF components and their combined weighted spatial distribution over time are presented in Fig 2.

**FIG. 2:**
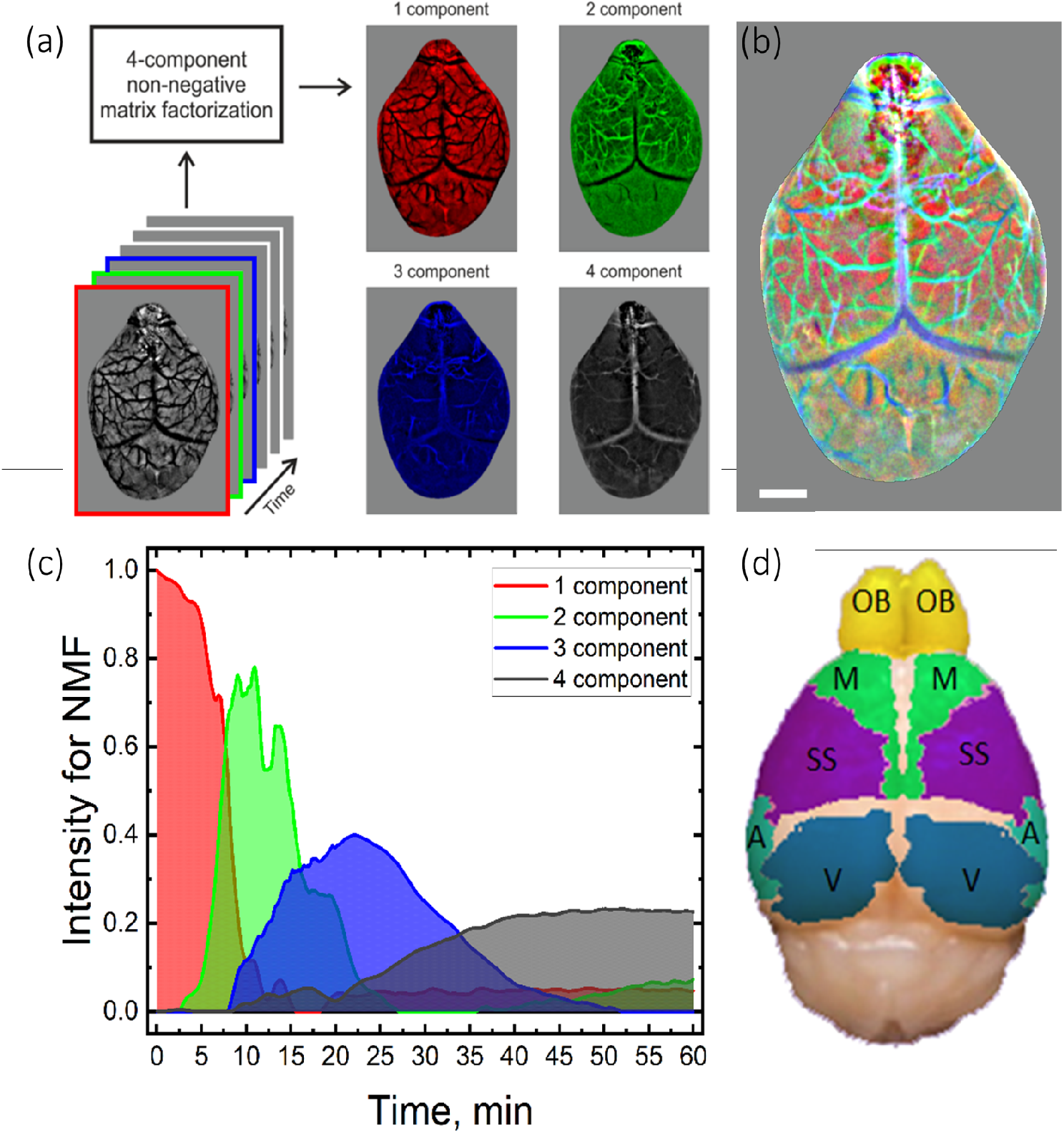
(a) Time series of LSCI images and corresponding color-coded images of the four NMF components (red, green, blue, and gray, respectively); (b) Merged image of the four color-coded NMF components. Supplementary video 1-d shows the dynamics of the spatial distribution of the merged four NMF components; (c) Weighted contribution of the NMF components in the merged distribution over time; (d) Corresponding map of the main areas of neuronal activity in the mouse brain: OB – Olfactory Bulb, M – Motor cortex, SS – Somatosensory cortex, V – Visual cortex, and A – Auditory cortex. Scale bar equals 1 mm.

The time series of LSCI images of the mouse brain obtained postmortem, as well as those processed using FFT, CWT, and NMF, show that the distribution of cerebral blood flow is highly non-homogeneous, with pronounced spatial and temporal hemodynamic localizations (see Fig 1, Fig 2, and associated Supplementary video 1). In line with previous studies that demonstrate a strong spatial relationship between hemodynamic changes and neural activity [28–30], also known as neurovascular coupling [31], the identified localizations of cerebral blood flow and blood microcirculation have been compared with a confirmed anatomical template and corresponding neuroanatomical domains [32]. The results obtained through LSCI, FFT, CWT, and NMF (see Fig 1, Fig 2, and Supplementary video 1) clearly reveal predominant hemodynamic localization in the olfactory bulbs. Additionally, successive alternate relocations of postmortem systolic blood micro-flow localizations, facilitated by the superior sagittal sinus, are observed between the somatosensory and visual cortical regions. Along with the results obtained by FFT and CWT (see Fig 1 and Supplementary video 1), NMF identifies temporal alterations of structural subcomponents, which exhibit a coherent evolution without being constrained by anatomical delineations (see Fig 2 and Supplementary video 1)

The selected NMF components are used to identify and evaluate the LSCI images in terms of their spatial variations in postmortem cerebral hemodynamics over time (see Fig 1, Fig 2, and Supplementary video 1). In the first 8 minutes following death, hemodynamic activity is predominantly observed in the cerebral cortex, and olfactory bulbs, visual, as well as in the cerebellum. The 1st NMF component corresponds to a rapid decrease in hemodynamic intensity during this period, followed by subsequent redistribution to the largest venous sinuses. The 2nd NMF component, appearing from the 4th to approximately the 25th minute, represents low-frequency pulsating blood flow in the sagittal sinus. The 3rd NMF component, active between 8 and 39 minutes, indicates further hemodynamic redistribution in specific cortical areas between arterial and venous vessels. Finally, the 4th NMF component, emerging around the 25th minute, is characterized by periodic hemodynamic relocations via the superior sagittal sinus.

The analysis of postmortem cerebral hemodynamics using LSCI, complemented by techniques such as NMF, FFT, and CWT, sheds light on the intricately complex process by which different parts of the brain cease to function following the cessation of blood circulation and oxygen supply. Although there is no definitive consensus yet, our observations contribute to the ongoing research in understanding this sequence. Preliminary findings suggest a potential order of impact: primary sensory areas, vital for basic sensory perceptions, are likely affected first due to their increased sensitivity to oxygen deprivation, as evidenced by the dominant hemodynamic activity observed in these areas immediately postmortem. This is followed by the impairment of higher-order sensory and motor areas, responsible for complex processing and coordination, which aligns with the redistribution of hemodynamic intensity detected in these regions. Association areas, crucial for higher cognitive functions, initially show resilience but gradually succumb to hypoxia, a pattern that can be traced through the temporal alterations identified by the NMF components. Ultimately, the frontal lobes, integral to executive functions, are among the last to be impacted, which correlates with the later stages of hemodynamic changes observed. It is important to recognize, however, that this sequence is not absolute and may vary across different species and under various conditions.

During the experiment, we observed a sequence of biological and physical phenomena related to brain activity and death, phenomena that have been extensively studied in prior research. Cardiac arrest, if not promptly compensated for, inevitably leads to the asphyxiation of all body organs, including the brain. Brain asphyxia lasting more than a few minutes typically results in brain death, a condition often used as a legal indicator of death. However, the definition of brain death is not universally accepted due to the variability in brain function cessation; different brain regions can cease functioning at different times. Brain death is frequently defined as the cessation of function in all parts of the brain, including the brain stem, yet cases exist where the brain stem remains active, enabling spontaneous breathing even when the rest of the brain has ceased functioning. In such cases, only life support equipment can maintain respiration. Distinguishing between brain death, coma, and chronic vegetative states presents a clinical challenge, as indicated by Bugge (2009) [33]. An absence of electrical activity, which can be detected through an EEG, may occur not only in brain death but also in deep anesthesia or during cardiac arrest.

Under conditions of oxygen deprivation, neurons lose their ability to generate action potentials, and glial cells also cease to function due to the failure of cell membranes to maintain the ionic balance between extra- and intracellular spaces. The resulting influx of ions causes a synchronous, massive depolarization of brain membranes, marking a transition from physiological to physical processes in the brain. Physically, this leads to brain swelling and compression of the capillaries, which can affect blood movement. This period is also characterized by a rapid transition from oxyhemoglobin to deoxyhemoglobin.

These intricate biological processes occurring during and after brain death can be precisely captured using optical brain imaging methods, including LSCI. It’s essential to recognize that the optical correlates observed through LSCI will depict significantly different processes before and after the onset of brain death. Currently, numerous questions about the biological processes associated with brain death remain unanswered. Notably, brain activity has been observed 10 minutes after the cessation of the heartbeat and the loss of all brain reflexes [4]. Additionally, experimental evidence suggests that conventional understanding of the rapid cessation of neuronal activity and irreversible changes in cellular structure post-brain death may need reconsideration. For instance, studies have shown that cellular functions of pig brain neurons can be restored hours after death. In one such study, four hours postmortem, a synthetic fluid called BrainEx, mimicking blood, was circulated through the pig brain [5]. This infusion of BrainEx was found to restore normal cellular metabolism, such as glucose utilization and ATP synthesis, in neurons and other brain cells, while preserving the structural integrity of these cells. Moreover, electrical stimulation led to the generation of action potentials in individual neurons [5].

However, the neuronal spikes observed were scattered and unsynchronized, indicating the absence of neural network functionality and, consequently, consciousness [34]. This finding aligns with research showing that mitochondria from the mammalian brain remain viable for several hours postmortem [35]. As a result, the idea that, under certain conditions, molecular and cellular functions in the mammalian brain might retain a partial capacity for recovery even after a prolonged postmortem interval appears reasonable [5]. Hence, current experimental and clinical data suggest that microcirculation and specific molecular and cellular functions in the brains of large mammals can be restored after brain death. It is increasingly evident that the molecular and cellular deterioration of the brain following circulatory arrest may be a more protracted process than previously believed, suggesting that the mammalian brain might possess a greater capacity for metabolic and neurophysiological stability under hypoxic conditions than is currently acknowledged [5]. While it may be somewhat premature to draw definitive conclusions about the process of neurovascular coupling in various brain regions during the terminal stage of hypoxia, the technique we have presented in this study marks a significant step forward in exploring this complex phenomenon.

These data will logically be compared with the clinical results of Emergency Preservation and Resuscitation (EPR) [36]. EPR technique was developed to use hypothermia to gain time during long-term cerebral hypoxia and allow delayed resuscitation. Also, animal studies have shown that cooling the body can cause circulatory arrest for up to two hours, followed by normal neurological recovery. This provides additional evidence that the process of irreversible destruction of neurons and astrocytes after cessation of blood supply to the brain may be longer than is commonly believed [36].

In conclusion, our pioneering application of NMF for both segmenting individual LSCI images and analyzing temporal series of these images has led to revealing significant temporal alterations in structural subcomponents. By employing a novel, integrated approach that combines LSCI, FFT, CWT, and NMF, we have succeeded in comprehensively analyzing cerebral blood flow dynamics in mice. This advancement not only contributes to our understanding of cerebral hemodynamics but also opens up new possibilities for exploring dynamic changes in brain function and pathology. This multifaceted methodology allowed us to metaphorically capture the ‘end of the tunnel’ experience, revealing intricate details of postmortem mice’s neural activity. The obtained results indicated a primary localization of hemodynamic activity in the ol-factory bulbs after death, suggesting a sustained sensory perception related to smell. This was followed by minor, but distinct, successive relocations of blood microflows between the somatosensory and visual cortical regions, facilitated by the superior sagittal sinus.

The adoption of this integrated approach has proven instrumental in elucidating the complex interplay of sensory processing and blood flow dynamics in the brain at the end of life. It provides a robust framework for interpreting the subtle shifts in neural activity and offers a comprehensive tool for visualizing and understanding these phenomena. This innovative methodological convergence opens new avenues for exploration into cerebral hemodynamics and sensory experiences postmortem. It holds the potential to bridge significant gaps in our neuroscientific understanding, particularly regarding the enigmatic aspects of consciousness and perception at life’s end. The insights gained from this study not only enhance our comprehension of cerebral physiology in mice but also lay the groundwork for future research that could extrapolate these findings to broader biological and philosophical contexts. An innovative artistic installation visually and conceptually illustrates what mice arguably perceive at the end of the tunnel (Fig 3). It combines the temporal variations of postmortem hemodynamic activities in the mouse brain with an artistic representation of this experience. The installation depicts the ‘end of the tunnel’ as a faintly flickering light in twilight, accompanied by the sensation of soft puffs of air and a strong ol-factory sensation, most likely akin to cheese. This artistic and scientific fusion in Figure 3 not only conveys our findings in a visually compelling manner but also provides a unique perspective on the sensory experiences of mice at life’s end. It opens new avenues for exploring these phenomena, potentially bridging the gap between neuroscientific understanding and the longstanding mysteries surrounding consciousness and perception at the end of life. The insights gained from this study enhance our understanding of cerebral physiology and sensory processing in mice, paving the way for further research into these complex and fascinating aspects of neural activity. In addition, the methodology developed in this study holds promise for broader applications in biomedical research, potentially paving the way for improved diagnostic tools and therapeutic strategies in neurology and various other medical disciplines.

**FIG. 3:**
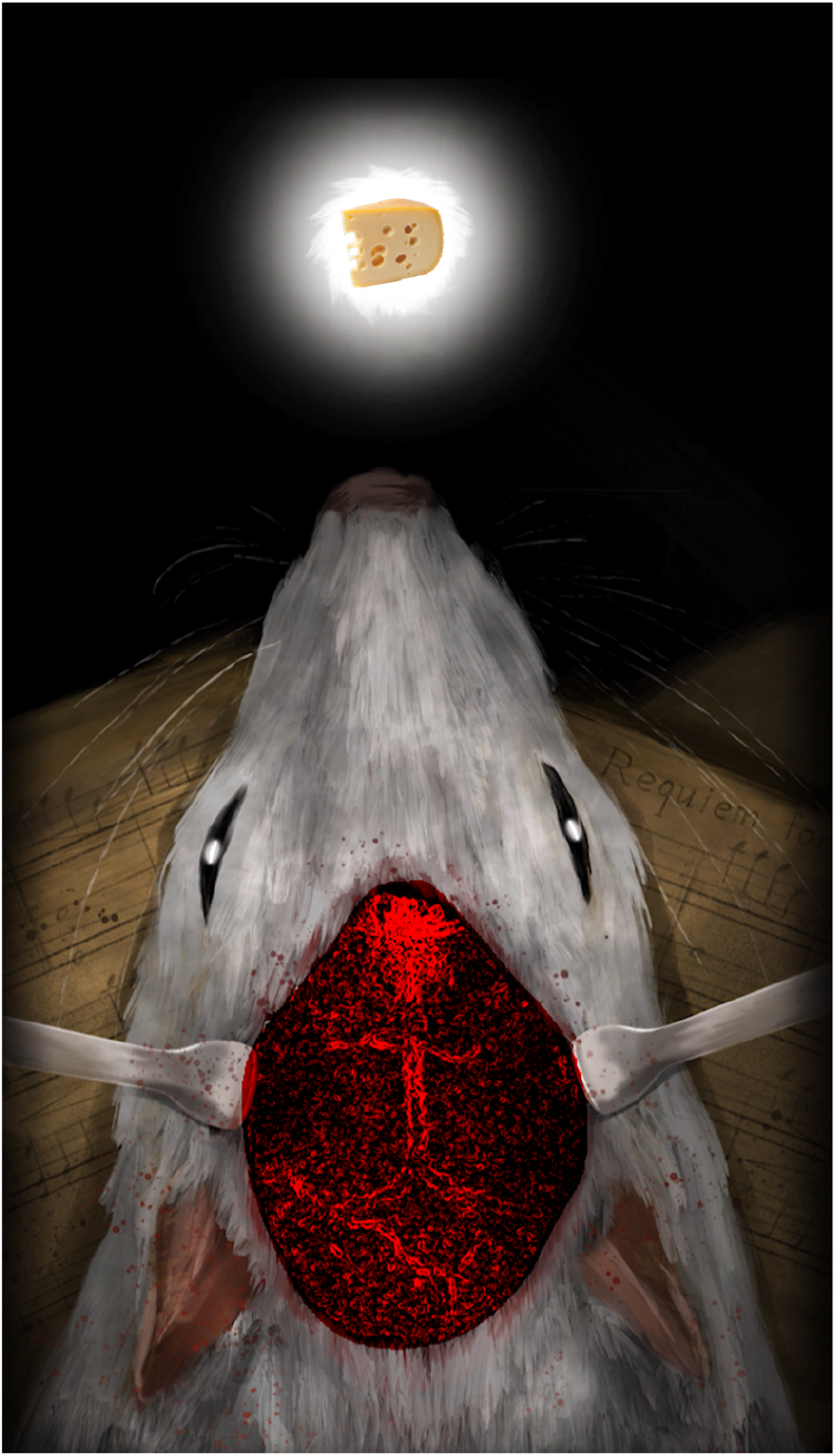
Artistic Interpretation of Mice’s Perceptual Experience at Life’s End. This illustration conceptualizes the ‘end of the tunnel’ vision for mice, as a blend of dim, flickering light reminiscent of twilight, coupled with sensations of gentle air currents and a distinct olfactory impression, likely resembling cheese. Supplementary video 2 complements this illustration by showcasing the temporal variations in postmortem hemodynamic activities within the mouse brain, integrated with this artistic representation of sensory experiences at the end of life.

## III. MATERIALS AND METHODS

### A. LSCI experimental setup

The developed Laser Speckle Contrast Imaging (LSCI) experimental system is a portable and effective non-invasive imaging technique. It is widely used for monitoring functional and morphological changes in blood perfusion, as well as relative blood flow variations in the brain [37], tumors [38], and other biological tissues in vivo [39, 40]. This setup has been employed for visualizing postmortem hemodynamics in the mouse brain. A 3 mW laser diode with an output wavelength of 808 nm (LDM808/3LJ, Roithner Lasertechnik GmbH, Austria) served as the light source. The beam from the laser diode passed through a diffuser (ED1-C20, Thorlabs, USA) to ensure uniform distribution of laser radiation over the observed brain area. Image acquisition was performed using a CMOS camera (DCC3240M, 1280 *×*1024 resolution, pixel size 6.7 *μm*, Thorlabs, USA) combined with a 12 mm F1.4 objective lens (Kenko Tokina Co., Ltd, Japan). The camera captured raw speckle patterns with an exposure time of 10 milliseconds. A series of 100 consecutive raw speckle images were obtained every 10 seconds during the experiment for further processing. Each series of images was then processed using a custom-developed temporal speckle contrast algorithm [5] based on MATLABr2019b software, assisted by a standard Fiji/ImageJ plugin [41]. This approach has been successfully used in transcranial vascular imaging [24] and cross-validated using fluorescent intravital microscopy [37], among other studies [42].

### B. Animal preparation

CD1 nude female mice, aged six to eight weeks and obtained from Envigo, were used in the study. All animal studies received approval from the Weizmann Institute of Science Institutional Animal Care and Use Committee (IACUC). The animals were anesthetized with a combination of 10 mg/100 mg/kg ketamine (Vetoquinol, Lure, France) and xylazine (Eurovet Animal Health, Bladel, The Netherlands) through intraperitoneal injection. Previous research has shown that the mouse brain’s vasculature can be clearly observed using LSCI through the intact skull. However, skin removal is still necessary for optimal visibility. Therefore, following the administration of general anesthetics, an initial incision was made, and the skin over the frontal, temporal, occipital, and parietal regions was removed by blunt dissection. This procedure was conducted to enhance the visibility of the brain’s blood vessels and minimize static scattering that affects ergodic conditions. The exposed area of interest was constantly moistened with saline. Subsequently, the animal was placed in the imaging setup on a specialized mouse holder, with measurements performed at room temperature (25°C). To register post-mortem brain hemodynamic activities, the animal was euthanized using a barbiturate overdose. The raw speckle images were captured within 60 minutes after the injection.

## Acknowledgements

Authors acknowledge help of Miss Tatian Koryashkina with the help of preparation of artistic installation presented in Fig 3.

## Author contributions

Conceptualization: IM; Methodology: IM, AS, VT, AB; Visualization: AS; Funding acquisition: IM, GP, AB; Project administration: IM, AB, GP; Supervision: IM, AB; Writing – original draft: IM, GP; Writing – review and editing: AB, GP, VT.

## Data and materials availability

All data is available in the main text or the supplementary materials.

## Methods

The methods were carried out in accordance with relevant guidelines and standard regulations, and are reported in accordance with the ARRIVE guidelines.

